# Temporal interference current stimulation in peripheral nerves

**DOI:** 10.1101/2022.07.20.500811

**Authors:** Ryan B. Budde, Michael T. Williams, Pedro P. Irazoqui

## Abstract

One strategy to electrically stimulate nerves utilizes the interference of multiple high frequency waveforms. This technique has recently gained significant attention as a method to improve the state-of-the-art in neurostimulation. Here we report our investigation into the fundamental properties of the neuronal response to these types of waveforms. Our data suggest, contrary to the currently accepted explanation, that neurons do not extract envelopes at all, and that the response to these signals is well explained by a resistor-capacitor (i.e., integrator) membrane with a fixed firing threshold. This new understanding of the fundamental mechanism of interferential neural stimulation changes how we should model and evaluate the safety and efficacy of these signals. Utilizing this new understanding, we develop several novel interferential stimulation techniques. Interferential strategies demonstrate promising results and may improve many neuromodulation therapies.

---

In 1948 Dr. Hans Nemec developed a novel neurostimulation technique based on the interference of two sinusoids^1^. This technique was used for decades with questionable efficacy in cosmetics and sports medicine, where it was called interferential current stimulation (ICS) or interferential current therapy (IFC)^2,3^. Some authors have previously explored the technique’s mechanisms^4^. In 2017 a group adapted the technique to a high performance, minimally invasive brain stimulator in a mouse model, which they called temporal interference stimulation (TIS)^5^. Since this initial publication the technique has garnered significant attention as a possible mechanism for noninvasive, precise, steerable brain stimulation^6–11^. We believe this technique will also bear great significance for peripheral nerve stimulators, which are useful in a variety of applications^12–19^. The technique utilizes the premise that 1) neurons extract envelopes, 2) neurons respond to different frequencies at different amplitudes, and 3) by utilizing a high frequency signal with a low frequency envelope we can create a stimulation field which is stronger in the middle, far from electrodes, than it is at the edges by the electrodes.

Previous work has not yet explored the fundamental relationship between interferential signals and the neuronal firing response in vivo. We explored these properties in experiments focused on the rat sciatic nerve. The accepted explanation for interferential neurostimulation, both at its inception and in recent work, is envelope extraction. Consider an example case where we stimulate a nerve with two sinusoids at 2000 Hz and 2020 Hz. At higher amplitudes either the 2000 Hz or the 2020 Hz signal would cause continuous neuronal activation, resulting in conduction block^20–23^. However, when these signals are applied together at a lower amplitude, we instead observe neuronal firing at 20 Hz (the beat frequency) – even though there is no 20 Hz energy in the system. In electrical engineering a beat frequency is extracted by rectifiers – a nonlinear circuit element which can generate 20 Hz energy from these two signals. The envelope extraction explanation for ICS assumes that neurons perform this function – that 20 Hz energy is generated – and that the neuronal firing is predominantly governed by this 20 Hz energy. With this explanation the most appropriate mathematical predictor for stimulation strength is the <u>minimum</u> of the two signals (i.e., the envelope amplitude), as both are required to produce the envelope. Our data suggest this explanation is incorrect, and that the neuronal response is instead governed by a passive resistor-capacitor membrane with a fixed voltage threshold, also called an integrate-and-fire model. With this explanation the most appropriate mathematical predictor is the <u>sum</u> of the two signals. With the integrator model the capability of the technique to produce stimulation fields that are stronger in the middle than they are on the edges is significantly reduced.

Here we use a variety of interferential techniques on the rat sciatic nerve to explore the ICS response. We apply up to 6 sinusoids and generate complex geometries and steering ability based on signals’ amplitude, phase, and interference properties. We modeled the fields generated and the neuronal response to them. Our data suggest that envelope extraction is a very minor contributor to the ICS response. Further, the Hodgkin-Huxley model poorly predicts the ICS response and integrate-and-fire models are better predictors. Our data suggest we should change the fundamental mathematics we use to simulate and predict the neuronal response to interferential signals, which will change conclusions – in some cases positively, in others negatively – about its efficacy, safety, and optimal applications. Our results suggest ICS may allow a clinically meaningful improvement in peripheral nerve stimulation.

Our new understanding of the mechanism will change the clinical use of ICS. Applications in which the target is small relative to the total stimulation area, such as noninvasive brain stimulation, are not as feasible as previously thought. However, precise control of stimulation regions near electrodes, like in cuff electrodes, is likely possible. By studying the ICS response, we might better understand the high-frequency biophysics governing voltage gated ion channels and update the Hodgkin-Huxley model.

## Results

We used a custom 12 contact cuff electrode placed on the rat sciatic nerve. We utilized a large number of unique stimulation parameters, and recorded the muscle responses in the biceps femoris and the plantar muscles. We then used neuronal and finite element models to predict and understand the responses we observed. Our experiments can be broadly grouped into two categories: exploration of the fundamental mechanisms and neuronal targeting. We report findings on 25 animals, with an additional 53 animals used in the development of the methods. A diagram of the cuff electrode and several types of waveforms are shown in Figure 1. Major findings are summarized in Table 1.

**Figure 1:**
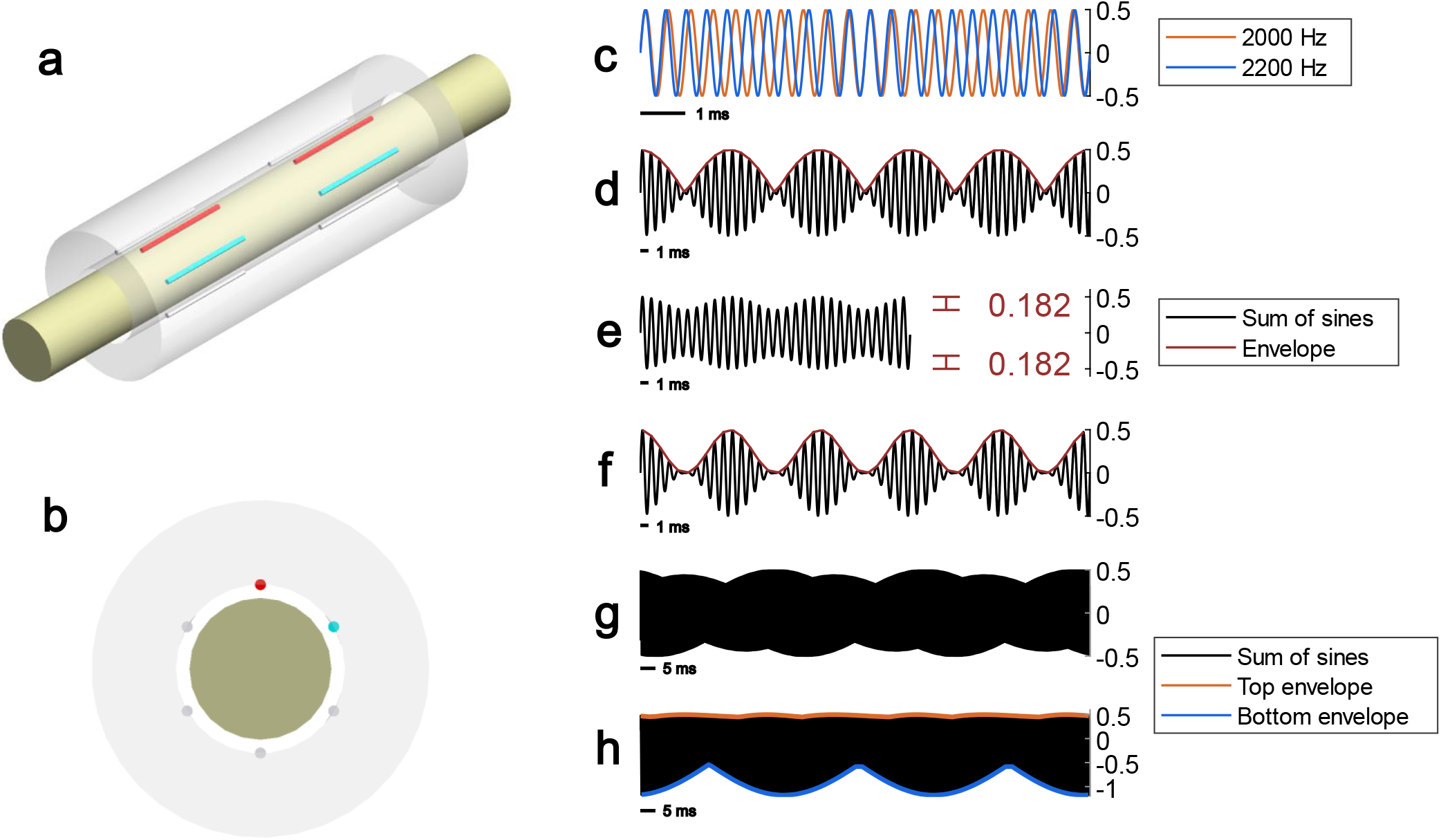
Tool and techniques used. a. CAD model of our cuff electrode. b. axial view of CAD model of cuff electrode. c. Example of interferential sines with 2000 Hz and 2200 Hz. These values are chosen to exaggerate the interferential effect. d. shows the sum of these two sines, with their envelope traced. The envelope of two sines is a rectified sine. e. Undermodulated 2 sine interference, which occurs when the amplitudes of the two sines are not equal. In this example the ratio is 1:4.5. f. 3 sine interference, which produces an envelope that is a pure sine. g. An example of secondary or higher ordered interference, with signals at 6000 Hz, 8000 Hz, and 10020 Hz. The amplitude of the envelope is greatly reduced. h. Our asymmetric interference pattern which consists of frequencies (Hz) – 2000, 2020, 4020, 6020, 8040; phase (radians) – 0, 0, π/4, π/2, π. Note the bottom envelope is much greater in peak-to-peak amplitude and distance from zero than the top envelope.

**Table 1:** comparisons regarding envelope extraction theory and an integrator-threshold theory

| Experimental result | n | Consistent with envelope theory? | Consistent with threshold theory? |
| --- | --- | --- | --- |
| Different envelope amplitudes and equal peak-to-peak amplitudes have identical recruitment | 4 | No. Envelope theory predicts identical recruitment at the reverse – equal envelope amplitude and unequal peak-to-peak amplitudes | Yes |
| Amplitude steering increases recruitment near the signal of increased amplitude | 2 | No. Envelope theory predicts the opposite – amplitude steering moves stimulation further away from the signal of increased amplitude | Yes |
| Increasing ICS amplitude transitions phasic firing to tetanic | 3 | No. Envelope theory predicts phasic firing and tetanic firing to have far-apart thresholds | Yes. Interference in space results in undermodulation, and once undermodulation crosses the activation threshold tetanus should be observed |
| Adding a second interferential sine to a tetanic single sine transitions tetanic firing to phasic | 2 | Indeterminate. Most envelope theory assumes phasic and tetanic thresholds are far apart, and their interplay is not discussed | Yes. Destructive interference reduces the tetanic sine below the firing threshold in a phasic manner, thus transitions firing activity from tetanic to phasic. |
| Firing threshold increases linearly with increasing frequency | 4 | Indeterminate. Hodgkin Huxley predicts a nonlinear increase. Ideal envelope extraction predicts no increase. Non-ideal rectification has been proposed. | Yes. Firing should increase linearly based on the resistor-capacitor values of the membrane. |
| Depolarizing envelopes drive neuronal firing | 4 | Indeterminate. Full-wave rectification is agnostic to hyperpolarizing vs depolarizing envelopes. Half-wave rectification only responds to one and not the other. | Yes. Neuronal activation is driven by a depolarized threshold, with the hyperpolarizing stimulus primarily necessary for charge balance. |

### Effects of envelope and carrier frequencies

In 4 animals we explored the effect of the envelope frequency (i.e., 5950 Hz and 6050 Hz compared to 5999 Hz and 6001 Hz) on the firing threshold. This effect has been previously modeled in^6,7^, who observed a “U” shaped response. We have also observed this response, as shown in Figure 3, albeit with a slightly different optimal frequency. We tested envelopes with a 6000 Hz carrier and a 2000 Hz carrier and did not see significant differences between data.

In 4 animals we explored the effect of the carrier frequency (i.e., 2000 Hz and 2020 Hz compared to 6000 Hz and 6020 Hz) on the firing threshold. We observed a linear response. In 2 of these animals, we additionally determined the threshold of single sinusoids (i.e., 2000 Hz and 6000 Hz on their own). <u>We found that the threshold was nearly identical for single sinusoids, which produce muscle tetanus and conduction block, and for interferential signals, which produce firing at the beat frequency.</u> In computer simulations we found that the Hodgkin-Huxley model predicts the response rises with frequency squared, not linearly, greatly overpredicts the thresholds, and at the amplitudes necessary to predict firing for interferential signals with a carrier over 4000 Hz we observed solver failure from floating point errors. In contrast, simpler models like the leaky integrate and fire (LIF) and generalized linear integrate and fire (GLIF) models predict a perfectly linear response and are much closer to real thresholds^24,25^. The computational models of neurons should always underpredict the firing threshold compared to real data because they use ideal currents, and our experiments suffer several nonidealities. Interestingly, the Hodgkin-Huxley model correctly predicts a linear frequency relationship for single sines^26^; it is only for spiking from 2-sine interference that the model breaks down. Even the GLIF model overpredicts firing thresholds at higher frequencies, suggesting its frequency-threshold slope is too high. We modified the GLIF model with a frequency dependent reduction in threshold, which can correct the slope with minimal impacts to lower frequency behavior. Frequency dependent capacitance might also correct these behaviors. Data are shown in Figure 2.

**Figure 2:**
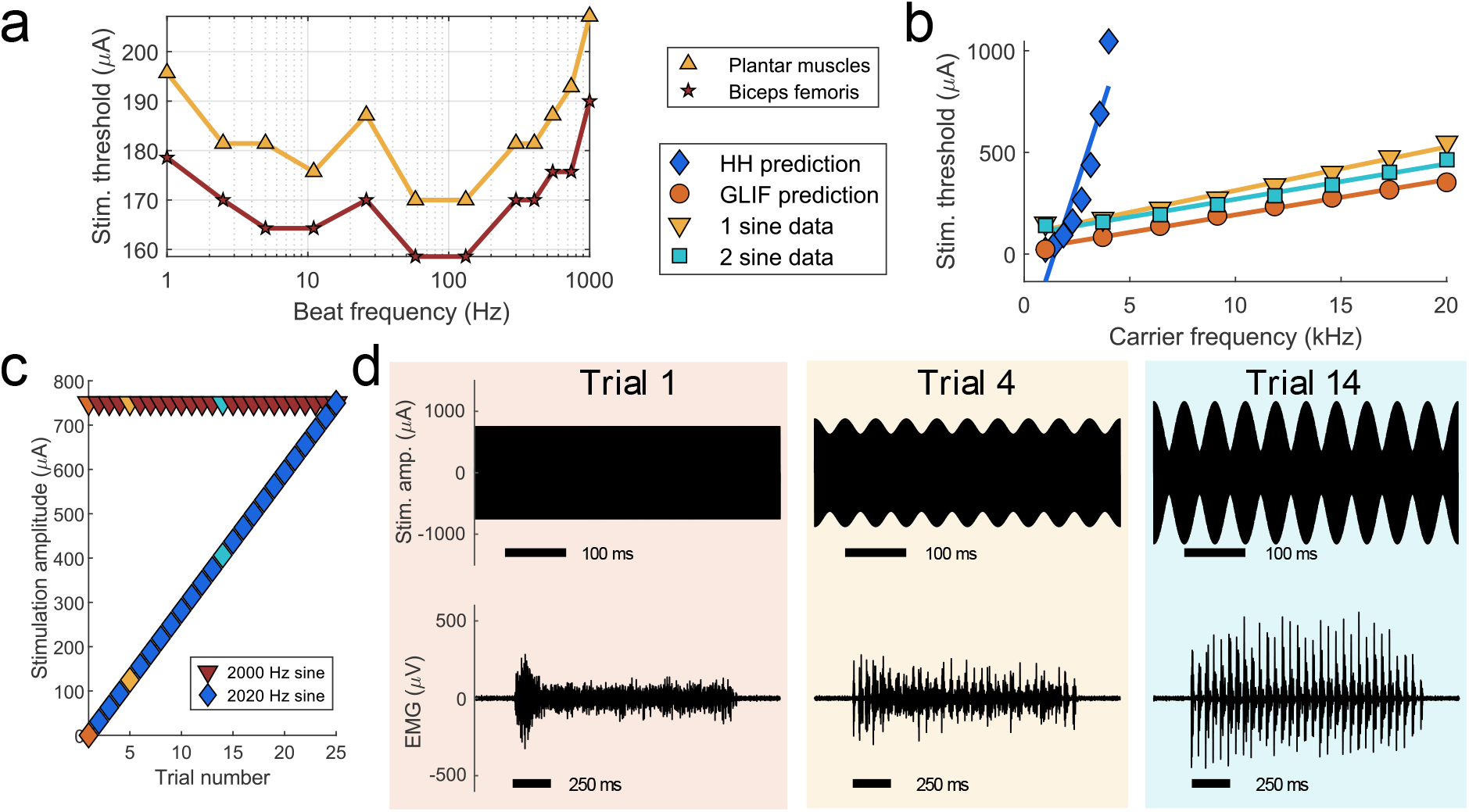
Envelope and carrier frequency effects; Interference transitions tetanic muscle activity to phasic. a. Animal data demonstrating the effect of the envelope frequency on the EMG threshold for a 6000 Hz carrier. Both muscles show a “U” shaped response, which has been predicted by simulations, albeit with different frequencies. b. Animal and neuron model data demonstrating the effect of carrier frequency on the firing threshold for a 20 Hz envelope. The 1 sine threshold, which produces tetanus, and the 2 sine threshold, which produces phasic firing, are very close and have a linear increase with frequency. The GLIF model is perfectly linear with frequency as it is an integrator. The Hodgkin-Huxley model is highly nonlinear when predicting the 2 sine response, despite the fact that it is linear when predicting the 1 sine conduction block threshold. Curve fitting suggests Hodgkin-Huxley rises with frequency squared. c. Stimulus amplitude of two sine interference. The 2000 Hz sine remains at constant amplitude and the 2020 Hz sine gradually increases in amplitude. Trials 1, 5, and 14 are highlighted. d. Data from selected trials. Top trace shown the stimulus. Bottom trace shows muscle activity. The stimulus begins as just a 2000 Hz sine, and transitions to a 2 sine interference pattern. EMG data begins tetanic and transitions to phasic. Data show strong phasic firing with inter-pulse amplitudes which are below the tetanic amplitudes, demonstrating destructive interference. The data suggest firing is dependent on peak-to-peak signal amplitude.

**Figure 3:**
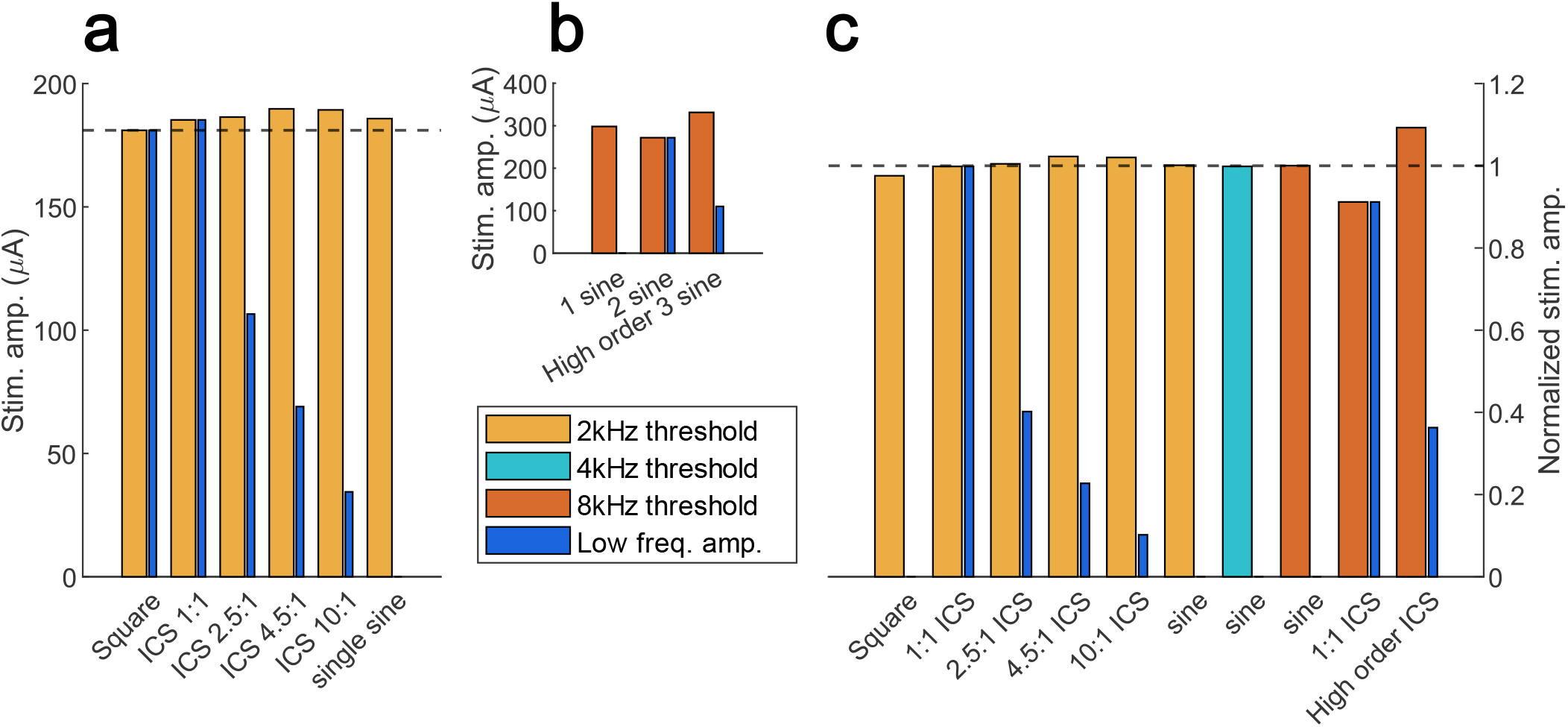
Trials with varying beat frequency amplitudes disprove envelope extraction as a major contributor to the ICS response. a. We used undermodulated ICS waveforms to investigate the effect of envelope extraction on the ICS firing threshold. In all cases the firing threshold was well predicted by the amplitude of the total signal, with the amplitude of the 20 Hz behavior playing a less significant or nonexistent role. Dotted line indicates the threshold of the square pulse. Signals: Square – 250 us pulse width 20 Hz repeat time, ICS – 2000 and 2020 Hz, single sine – 2000 Hz b. We use ICS signals in which the 20 Hz behavior was either a primary (2 sine) or secondary (3 sine) interference. In these strategies the amplitude of the 20 Hz behavior is very different, but the firing threshold were similar. Signals: 1 sine – 8000 Hz, 2 sine – 8000 and 8020 Hz, 3 sine – 6000, 8000, 10020 Hz. c) Using the frequency linearity of signals we can normalize the data from plots a and b. All thresholds are similar once their frequency response has been accounted for. Dotted line is at 1, the normalized predicted firing threshold. Signals – as described for a,b, and an additional 4000 Hz single sine. The data suggest firing is independent of low frequency behavior.

### Undermodulation results contradict envelope extraction theory

Based on our results regarding the effects of the carrier frequency we hypothesized that neurons may not extract envelopes. To test this theory, we devised multiple strategies which maintained the same peak-to-peak amplitude for a given signal but had different amounts of envelope amplitude, through a technique called undermodulation (See Figure 1). Consider signals at 2000 Hz at 100 µA and 2020 Hz at 100 µA compared to signals at 2000 Hz at 180 µA and 2020 Hz at 20 µA. The 20 Hz behavior in the second set is much smaller, with an envelope peak-to-trough value of 40 µA compared to the 200 µA in the former case, with the same peak-to-peak amplitude for both cases. Therefore, if the first set of signals produces firing, then envelope extraction theory predicts the second set will not, while a fixed threshold theory predicts it will. We also utilized higher order interference signals to test this theory (see Figure 1). Consider signals at 8000 Hz at 100 µA and 8020 Hz at 100 µA, compared to signals at 6000 Hz at 66 µA, 8000 Hz at 66 µA, and 10020 Hz at 66 µA. The amount of 20 Hz behavior in the second set is again much smaller. For both undermodulation and higher order interference we observed firing thresholds consistent with a fixed threshold theory, not an envelope extraction theory. We verified these effects in 2 animals, and an additional 2 animals in which we only compared the 1-sine-tetanic-activity and 2-sine-phasic-activity thresholds. These data are shown in Figure 3.

### Responses to multiple types of interference

In 5 animals we explored the firing behavior in response to signals with “secondary interference.” Consider four signals at 3000 Hz, 5000 Hz, 12000 Hz, and 14020 Hz. The 3000/5000 group produces an interference pattern at 2000 Hz, and the 12000/14020 group produces interference at 2020 Hz. The sum of these patterns interferes at 20 Hz. We also used a technique with signals at 6000 Hz, 8000 Hz and 10020 Hz (Shown in Figure 1). We observed muscle activation at the beat frequency of 20 Hz for these techniques. The additional “layer” of interference in these signals means the amount of 20 Hz behavior is much smaller than the peak-to-peak amplitude. The firing thresholds for these techniques were consistent with the peak-to-peak amplitude, not the 20 Hz behavior, supportive of a fixed-threshold mechanism.

In 5 animals we explored how constructive and destructive interference between signals can alter the response. In 3 animals we applied a single sine at 2000 Hz and 0 degrees of phase at a constant amplitude sufficient to cause muscle tetanus. We then gradually increased the amplitude of a second signal at 2000 Hz and 180 degrees in phase. These signals destructively interfered in space, causing muscle activation to cease. In 2 animals we applied a single sine at 2000 Hz at a constant amplitude which caused muscle tetanus. We then gradually increased the amplitude of a second signal at 2020 Hz. As the amplitude of the 2020 Hz signal increased the tetanus disappeared and was replaced by phasic firing at 20 Hz, as these two signals sometimes constructively interfere, to produce muscle activation, and at other times destructively interfere, producing silence. The reduction in motor activity during the periods of destructive interference is supportive of a fixed-threshold mechanism. These data are shown in Figure 2.

In 1 animal we utilized a technique proposed by Cao and Grover^6^, which uses six sinusoids at 2000 Hz, 2005 Hz, 2010 Hz, etc. up to 2025 Hz. We started with 2000 Hz and 2005 Hz, and then added 2010 Hz, and then 2015 Hz etc. We observed that in each case the firing threshold is consistent with the total current input. We predict this strategy will be useful for optimal ICS strategies as it maximizes the interference amplitude while minimizing the amplitude of each contributing signal.

In 3 animals we attempted 3-sine interference utilizing signals of 1980 Hz, 2000 Hz, and 2020 Hz, where the 2000 Hz signal was twice the amplitude of the others, as compared 2-sine interference utilizing just 2000 Hz and 2020 Hz of equal amplitudes (see Figure 1). Three-sine interference produces an interference pattern that is a pure sine, compared to a rectified sine pattern produced by 2-sine interference. We observed no significant improvements.

### Blocking behavior

Others have predicted that higher amplitude ICS signals can cause conduction block^7^. In 3 animals we increased the stimulation amplitude of ICS signals well beyond what was necessary to elicit motor contractions. As stimulation amplitude increased, we observed a gradual decrease in motor activity until we only observed onset and offset firing, consistent with conduction block. In 1 animal we also saw an increase in activity again, as we hypothesize that we started to activate an adjacent neuronal track which innervated the same muscle. These data are shown in Figure 4.

**Figure 4:**
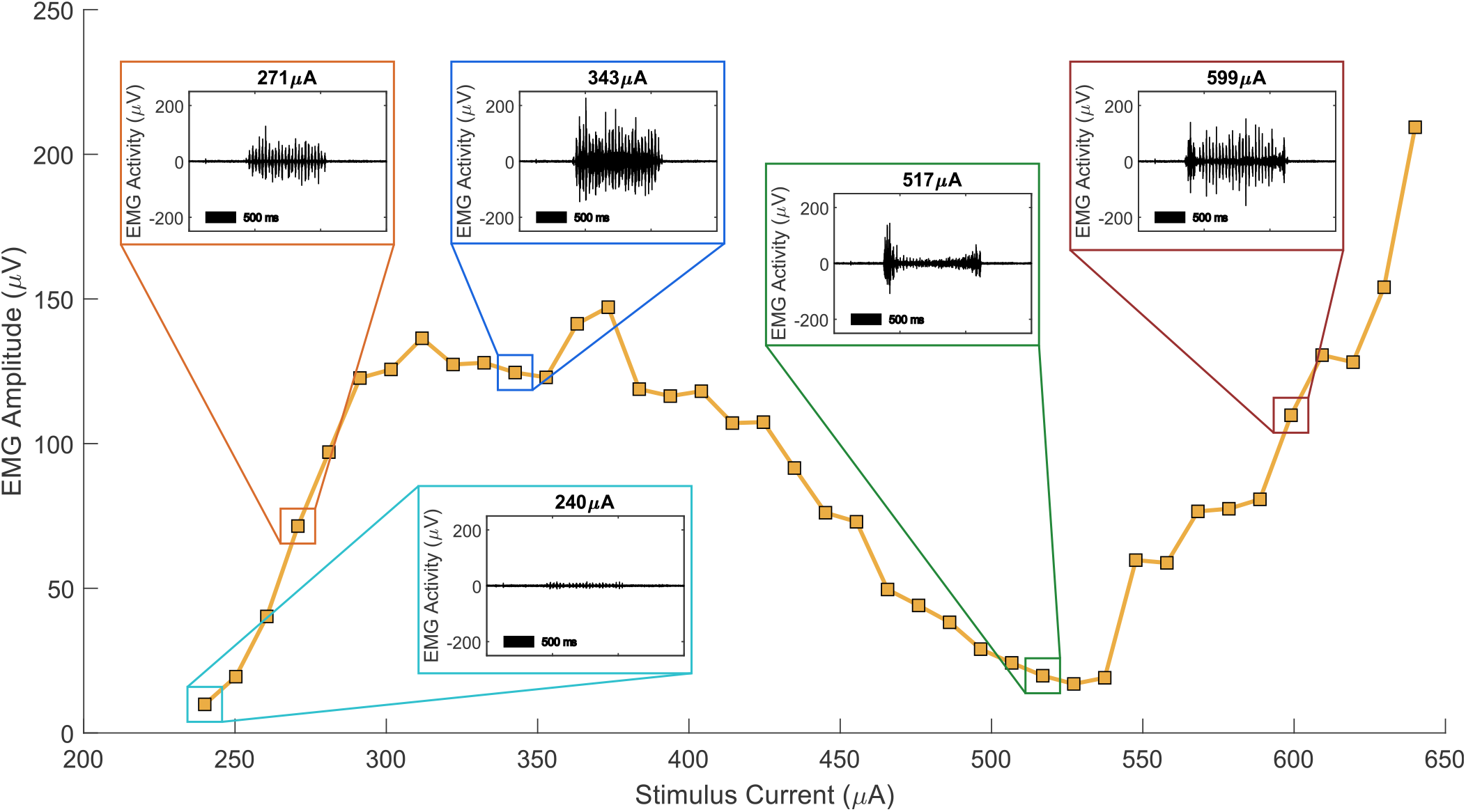
Interferential signals can transition from phasic firing to conduction block at high amplitudes. The primary curve shows the peak EMG activity for each trial. The insets show the raw EMG data for each trial. These trials use a 2 sine interferential stimulus with sines at 2000 Hz and 2020 Hz. As the stimulus amplitude increases the EMG activity reduces, with onset and offset activity. After sufficient amplitude the onset and offset remain, but new phasic firing appears, which we hypothesize is caused by activation of a different nerve innervating the same muscle group.

### Geometric targeting and mapping

Until now we have discussed experiments which focus on the fundamental response to interferential signals and our conclusions apply generally and are not dependent on the geometry of the field created. Next, we will discuss experiments in which the geometry impacts our conclusions. We utilized our 12-contact cuff electrode and multiple types of signals to generate geometrically complex regions of stimulation. We utilized one geometry, our mapping geometry, which targeted a 60-degree slice of the nerve near the edge of the cuff. By rotating this geometry around the cuff across 6 trials we were often able to obtain good separation between our two muscle targets. Using these data, we generated an estimated map of each target within the nerve, and then utilized this map to inform subsequent trials. An example of the mapping process is shown in Figure 5. In a series of experiments, we used two sine interference (n=2), three sine interference (n=3), and higher ordered interference (n=4) along with complex geometries to change muscle recruitment. We used phase interference (n=3) without changing the geometry to similarly target muscle recruitment. In these phase experiments we used three signals: 2000 Hz at 0 degrees, 2020 Hz at 0 degrees, and 2000 Hz at a variable phase (0 to 180 degrees). Because the signals are all generated from different contacts there are multiple regions of constructive and destructive interference, allowing the generation of both stimulation “hot spots” and “cold spots.” Importantly, phase interference allows us to effect a greater change on the field at a distance from the electrodes than at the electrodes, and is, so far, our only technique to do so. All techniques were able to significantly change the relative recruitment of muscles. Data are shown in Figure 6. In Figure 6e a difference in phase causes an increase in biceps femoris activation even though there is no increase in field strength anywhere in the region as a result of this phase change (there is a reduction in activity in many places, explaining the reduction in plantar muscle activity), suggesting activation is not determined solely by field amplitude.

**Figure 5:**
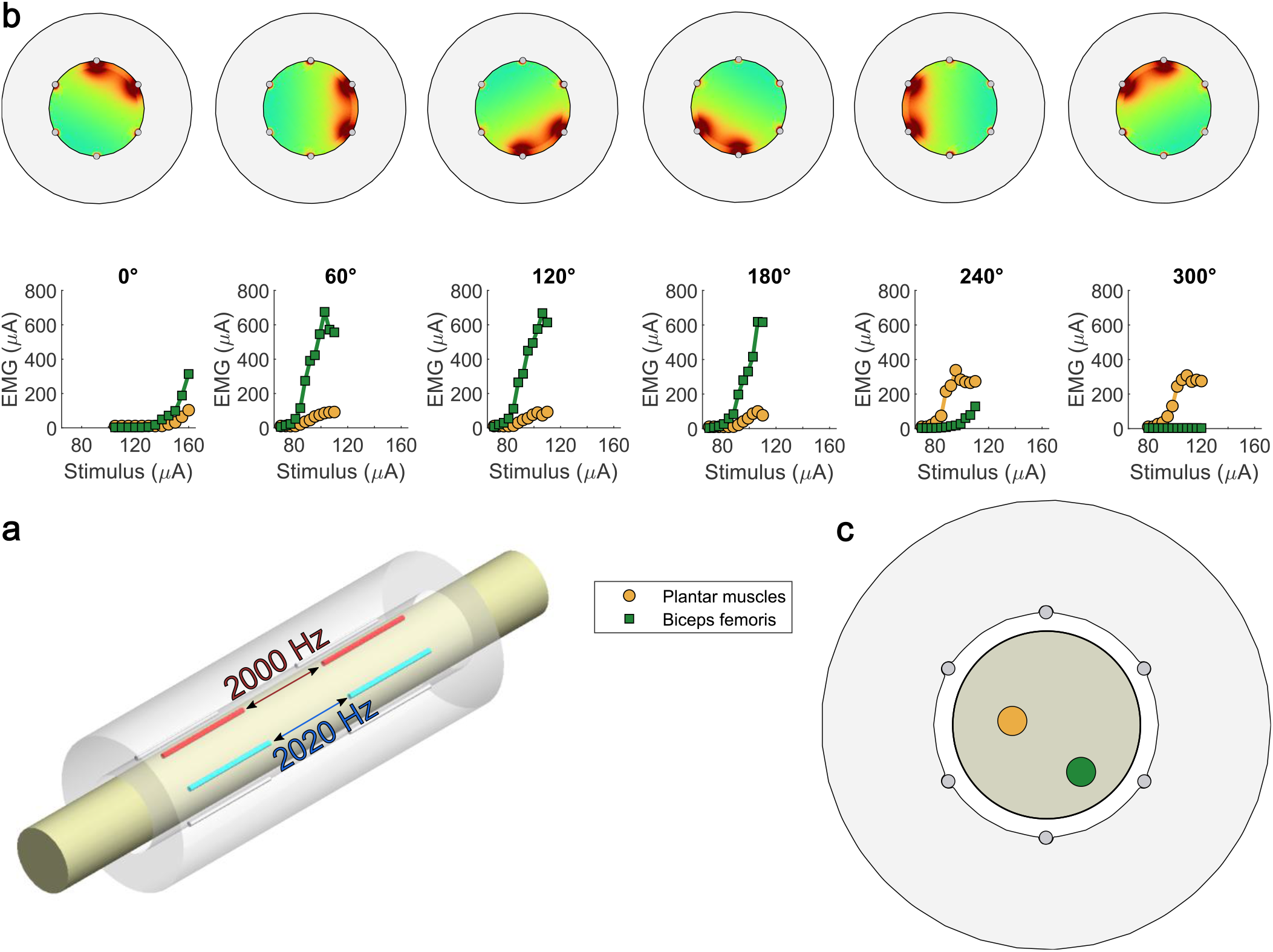
Mapping process. a. The stimulation geometry used for mapping. This geometry creates a tight stimulation region along a 60-degree arc of the nerve and usually allowed excellent muscle selectivity. b. Mapping data. With our cuff electrode we could rotate this stimulation geometry six times around the nerve. Top images show the electric field strength for a given rotation, and bottom traces show the recruitment curve data. c. The predicted nerve branch locations. The predicted locations are consistent with the observed data. Both muscles require high amplitudes in the 0° rotation, and so their locations are further from this region. The biceps femoris is preferentially activated in rotations 60°, 120°, and 180°, and so its predicted location is near these regions. The plantar muscles are preferentially activated in rotations 240° and 300°, and so its predicted location is near those regions.

**Figure 6:**
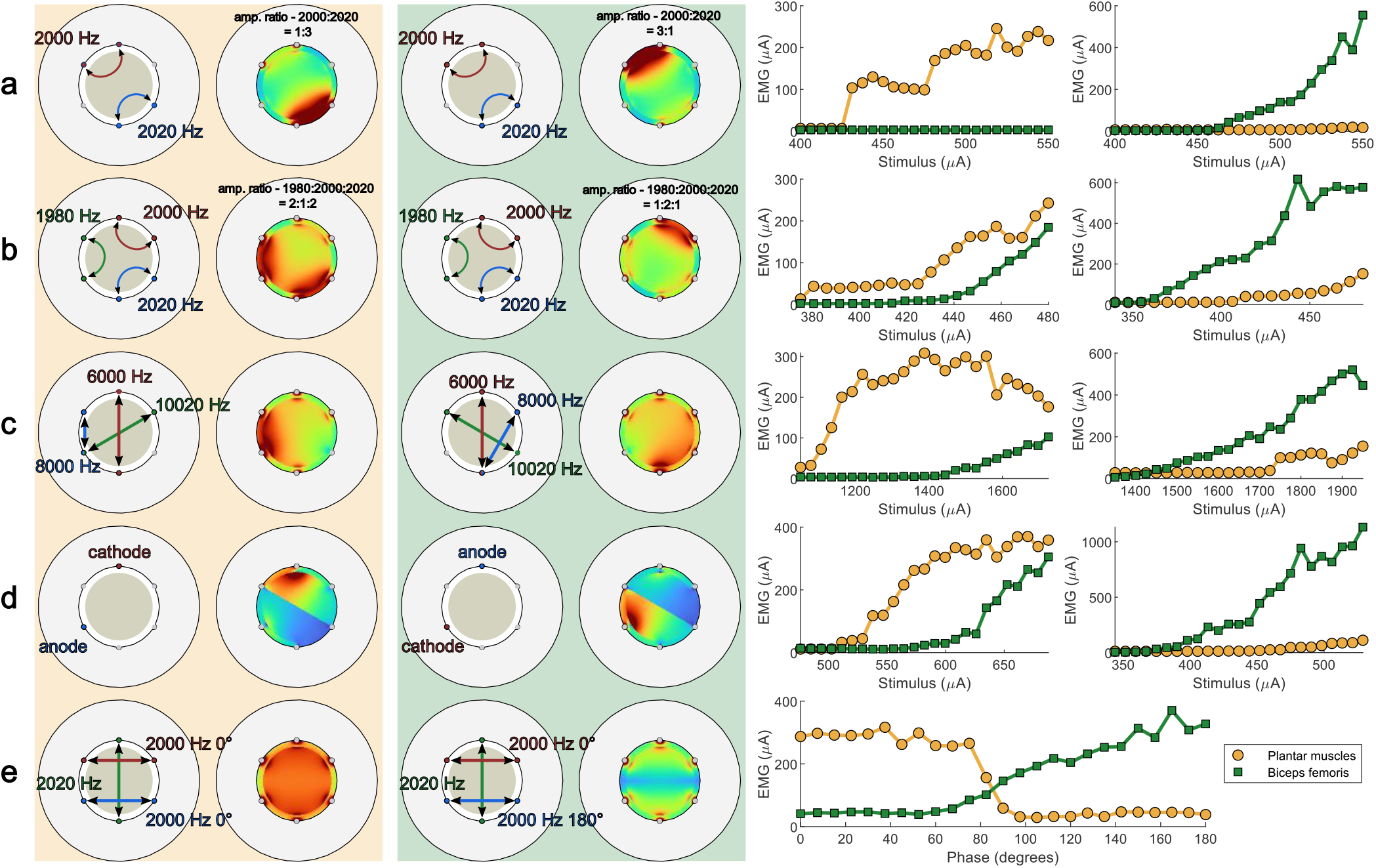
Muscle selectivity achieved through various techniques in multiple animals. Each row shows, from left to right: connections for plantar-preferred geometry, electric field strength for plantar-preferred geometry, connections for biceps femoris-preferred geometry, electric field strength for biceps femoris-preferred geometry, data for plantar-preferred geometry, data for biceps femoris-preferred geometry. a. Amplitude steering with 2-sine interference. b. Amplitude steering with 3-sine interference. c. Different high-order interference geometries. d. Asymmetric envelope, where selectivity is caused only by inverting the signal. e. Phase steering, where selectivity is caused only by changing the phase of one signal. This trial only requires one curve to observe selectivity, where the amplitude for all signals remains constant, and instead the phase of one signal is gradually changed.

### Asymmetric envelopes

Neurons should fire more readily in response to a depolarizing signal than a hyperpolarizing signal of similar amplitudes. We hypothesized this property would extend to interferential signals. We devised a charge-balanced waveform that had very different depolarizing and hyperpolarizing peaks (See Figure 1). We then applied this signal and its inverse and observed significant differences in motor recruitment in 4 animals. When we model this effect, we observe a sharp discontinuity because we are only interested in the most depolarized potential for each location in space. These data are shown in Figure 6.

### Amplitude steering results contradict envelope extraction theory

One benefit of ICS is that the stimulation field is steerable – the targeted neuronal region can be moved because the region is created by multiple signals originating from different points in space. The most proposed method of steering is changing the relative amplitudes of the signals. For example, if we are using a 2000 Hz signal on the left side of a nerve and a 2020 Hz signal on the right side of the nerve, then activation should be symmetric about the center of a nerve. According to envelope-extraction theory, if we increase the amplitude of the 2020 Hz signal, then the region of activation will move towards the 2000 Hz signal, since activation is predicted by the minimum of the signals. However, with a fixed-threshold theory an increase in the amplitude of the 2020 Hz signal will move the region of maximal activation toward the 2020 Hz signal. In 2 animals we first obtained multiple measurements that allowed us to estimate the location of each motor group within the nerve. Next, we used two sines to cause activation, and increased the amplitude of one or the other sine in order to steer the signal. In both animals steering causes an increase in motor activity in the motor group that was closer to the signal that was increased. These data are shown in Figure 7. These results are consistent with a fixed-threshold theory and contradict an envelope-extraction theory. We continued to change the relative ratios of these two sines until we only stimulated with a single sine. By envelope extraction theory the stimulus region should disappear, and we would see a sharp difference in motor activity, while a fixed-threshold theory predicts little to no change in activity once the signal is fully steered to only a single sine. We observed no significant change when moving from a strong steer to a single sine. We also observed motor activity transitions from phasic to tonic, suggesting motor activation is primarily driven by the high amplitude single sine.

**Figure 7:**
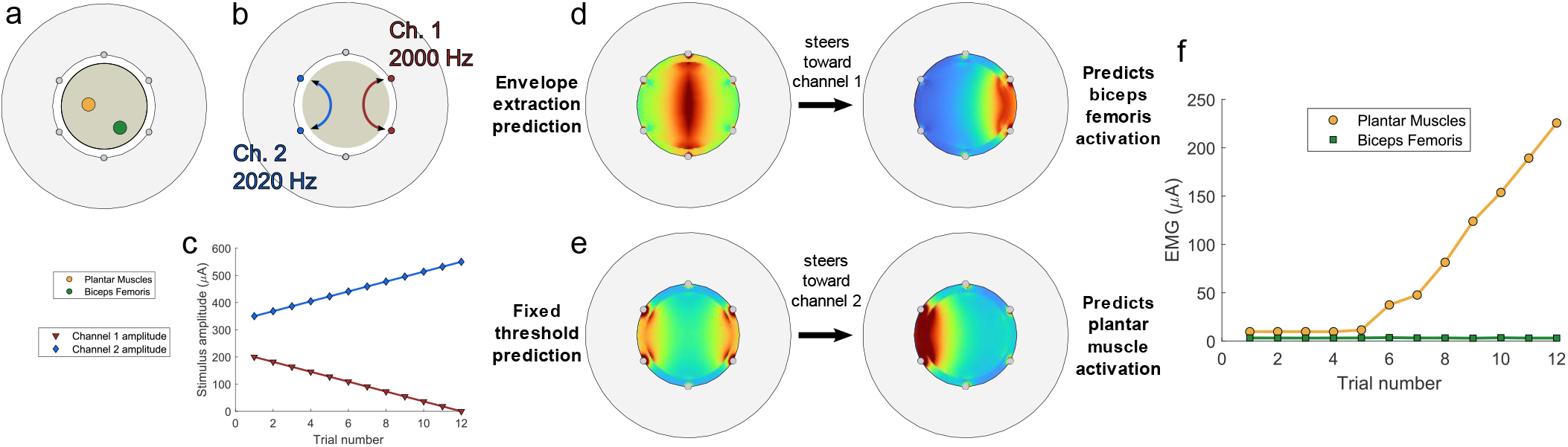
Amplitude steering predictions. a. The predicted map of sub-branch locations for this animal. b. The stimulus diagram for this experiment. The first channel is on the right side, nearest the biceps femoris sub-branch, and the second channel is on the left, nearest the plantar muscle sub-branch. c. Stimulus parameters. In this experiment the interferential focus is gradually “steered” by increasing the amplitude of Channel 2 and decreasing the amplitude of Channel 1. The total current amplitude remains constant. d. Envelope extraction theory predicts the stimulation region will move toward the right, activating the biceps femoris. e. Fixed threshold theory predicts the stimulation region will move to the left, activating the plantar muscles. f. Results. The plantar muscles are strongly activated, supporting fixed threshold theory and contradicting envelope extraction theory.

## Discussion

Our results suggest that the neuronal response to interferential stimulation is well predicted by an integrator model and poorly predicted by an envelope extraction model. The data suggest the Hodgkin-Huxley model, while accurate for single high frequency sines, does not accurately predict firing in response to two high frequency interferential sines. These conclusions suggest that interferential techniques as originally proposed – to target a small brain region used amplitude-based steering without significant side effects – may not be feasible because stimulation regions are not stronger in the center than they are on the edges, at least with published techniques. However, by utilizing the strategies we discuss or adapting to new targets we predict that interferential techniques will still be able improve efficacy in a range of neurostimulation applications.

We explored a variety of interferential techniques - including construction, destruction, phase differences, asymmetry, and secondary or tertiary interferences, and they can all cause differences in recruitment. These techniques, in addition to geometric current steering based on a multi-contact cuff electrode, may enable greatly improved neuronal targeting in peripheral nerve stimulators. Such specificity may improve efficacy and reduce side effects in clinical applications. Further, such targeting may allow a transition towards less restrictive cuffs, which may also improve patient outcomes. The novel techniques we propose demonstrate targeting ability which expands upon previous techniques. Combinations of complex targeting techniques may eventually deliver on the promise of ICS and enable minimally invasive brain stimulation.

Classical neuronal models predict neuronal firing as primarily dependent on the derivative of the field in the axis of the nerve^27^. However, in these models the stimulus only was applied in the axis of the nerve, and so derivatives in other directions were already zero. Our data suggest the activation function may be isotropic, and any derivative of the field may drive firing. Some interferential techniques may be able to cause significant changes in the derivative of the field in the center of the target (i.e., phase destructive interference), better enabling minimally invasive stimulation.

### Outlook

Further work must be done to establish the single axon response to interferential signals and construct an accurate, biofidelic model. The use of interferential signals may allow new insights into fundamental neuronal physiology.

So far, we have only tested each interferential technique in isolation. Interferential neurostimulation is a promising technology because the complexity of these techniques is multiplicative. We can combine complex geometry via the 12 contacts <u>and</u> phase induced hot and cold spots <u>and</u> higher order interference patterns <u>and</u> asymmetrical interference. These techniques are also only targeting neurons by their physical location. By optimizing the envelope frequency, we might also target neurons based on their diameter, a technique already used in pulse-width optimization of fiber type recruitment^28^. We also hypothesize that it is possible to generate standing waves in tissue, which could be used to generate even more complex stimulation patterns. When these techniques are combined the stimulation complexity for a single patient may be too great to reasonably explore without learning algorithms.

## Methods

### 12-contact peripheral cuff electrode

We utilized drawn filled tube (DFT) wire with a 90% platinum 10% iridium alloy outer sheath (20% of total mass) with a tantalum core (80% of total mass), with a wire diameter of 50 μm and PFA insulation to a total outer diameter of 75 μm (Fort Wayne Metals). We sewed the wire into a cut split silicone tube (ID 0.04” OD 0.085”, AM systems). The cuff was 4 mm long, with two rings of contacts, each 1 mm in length with 1 mm spacing. Each ring contained 6 evenly radially spaced contacts which ran parallel to the nerve. The insulation was removed to create exposed electrode contacts inside the cuff. A rendering of the cuff on the nerve is shown in Figure 1.

### Stimulator circuit

A full circuit schematic is shown in the supplement. We utilized a mirrored Howland current source as proposed by^29,30^ for each stimulator channel. We additionally included a DC blocking capacitor at the output, to prevent DC leakage during stimulation, which can skew recruitment curves^31^. We utilized MIC920 operational amplifier and 0.1% tolerance resistors to keep the output impedance high. We used DG419 switches to discharge the DC blocking capacitors after each trial. The input of each stimulator was driven by an analog output channel on a National Instruments USB 6343 Data Acquisition unit. Output currents were validated by applying the currents across known resistors and measuring the voltage.

### Neuronal models

We utilized single axon patch neuronal models from the Hodgkin Huxley and GLIF models^24,25,32^. We used the default parameters from each paper. For the GLIF model we used the parameters for characteristic neuron L, for hyperpolarization bursting. We additionally modified the threshold term for the GLIF model to correct its slope at high frequencies. While the original equation is given by:

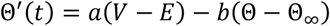

We instead used

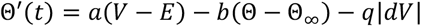

Where q is small relative to a or b. This change lowers the threshold for firing in the presence of a strong, high frequency stimulus, with minimal impact on low frequency behavior. We used a value of q = 1.28e-4, with a = 30 and b = 10. For each trial we simulated 500 ms with stimulation applied from 100 ms to 400 ms. We used carrier frequencies from 1000 Hz to 20,000 Hz and a beat frequency of 20 Hz. We included a pi/2 phase offset so the stimulation period began with the beat amplitude at zero, to prevent edge effects. A stimulation was deemed above threshold once there was at least one spike per beat in the time range 150 ms – 350 ms. The GLIF model was solved with the finite difference method with steps of 10 µs, and the Hodgkin Huxley model was solved with MATLAB’s stiff ODE solver ode15s. After the Hodgkin Huxley model displayed instability we adapted it to a compartment model, to see if spikes would propagate outside of the initial patch, but they did not. Thresholds were calculated to a resolution of 1 µA/cm^2^.

### Electric field models

We modeled the fields produced by various geometries using COMSOL Multiphysics 6.0. We used a steady currents model as recommended by^33^. We used constant potential boundary conditions, with each positive channel source modeled as 1 V and each negative channel source modeled at -1 V. Each channel was modeled as completely independent and were superimposed for analysis. We used impedances based on cerebrospinal fluid for the fluid bulk, and biofidelic impedances for the other materials. All impedances were based on literature values measured at 1 kHz. We exported the data into MATLAB 2021b for further processing and analysis. For our setup a negative potential is depolarizing. All our model images calculate the most negative potential for each voxel, and then calculates the magnitude of the electric field for that voxel for that moment in time. Therefore, our images do not represent a snapshot in time but rather a rough estimate of firing probability over the course of the entire stimulation period. We chose not to use the activation function as it is a metric which follows from stimulation which is in the axis of the nerve and where Hodgkin Huxley predicts firing. Given our stimulation is frequently not in the axis of the nerve, and given our data suggests Hodgkin Huxley does not accurately predict firing, the activation function is not the most appropriate metric.

### Surgical procedures

We present data on a total of 25 animals, with an additional 53 animals used in the development of the methods. All animal procedures were approved by the animal care and use committee of Johns Hopkins University. We utilized both male (n = 13, weight = 348 g ± 75.2 g) and female (n = 12, weight = 257 g ± 22.0 g) Long Evans rats. Animals were anesthetized with urethane (1.5 g/kg i.p.), shaved, and body temperature was maintained at 37 °C with a thermostatically controlled heating blanket (Harvard Apparatus). The choice of leg (left or right) was determined by a coin flip. We placed the animal prone and oriented the hip and knee at 90 degrees. We made an incision along the skin 4 mm distal and parallel to the femur. We blunt dissected the surrounding tissue to expose the biceps femoris, which we reflected caudally to expose the sciatic nerve. We followed the sciatic nerve from the hip to the trifurcation into the tibial, sural, and peroneal nerves. We gently blunt dissected connective tissue and placed the cuff electrode just proximal to the trifurcation. We carefully tied two sutures around the cuff to keep it closed, placed at the edges of the silicone. We filled the surgical pocket with lactated Ringer’s solution and covered the wound with plastic wrap to prevent it from drying out. We next expanded the subcutaneous pocket to expose the surface of the biceps femoris. We placed two 30G stainless steel needles bent into hook electrodes, placed approximately 14 mm apart, 6 mm distal and parallel to the femur, with each hook penetrating tissue approximately 2 mm wide and 2 mm deep. We placed two similar hook electrodes in the plantar of the foot, one near the ankle and one near the fifth digit. The hind limb was placed on a rigid plastic board with the animal prone, the knee and hip at 90 degrees, the ankle at 0 degrees, such that the foot was fully plantar flexed with the plantar facing upwards. We placed tape across the knee, ankle, and toes, just outside the recording region, in order to keep the leg in place and reduce motion artifact. We placed a stainless steel 1 inch ground plate on the skin near the base of the tail using conductive gel (Tensive).

### Stimulation trials and parameters

Unless otherwise stated, each stimulation trial consisted of a 0.5 second ramp up, 1 second of full amplitude stimulation, and a 0.5 sec ramp down, to prevent onset and offset effects. Waveforms were generated and muscle signals recorded in MATLAB at 250,000 samples per second via a National Instruments USB 6343 multipurpose data acquisition system. Data was recorded for an additional 0.5 second before and after each stimulation trial. In all interferential techniques we used pure sinusoids generated by code in the range 1980-20,020 Hz. Unless otherwise stated, most trials were performed with a carrier of 2000 Hz and an envelope of 20 Hz. We utilized some traditional bipolar waveforms as a point of comparison. Bipolar square cathodal pulse repeat frequency was identical to the envelope frequency, usually 20 Hz, with a comparable pulse width, 250 µs pulse width (500 µs period) being equivalent to a 2 kHz sine. In trials with a very slow beat frequency (i.e., 1 Hz) we extended the full amplitude stimulus duration to be 20 periods.

We began each experiment by creating a list of desired stimulation techniques. We next randomized the order of these techniques. For each technique we first tested individual parameters in order to find the upper and lower stimulation thresholds for muscle recruitment. Next, we ran a recruitment curve program which would automatically generate evenly spaced amplitudes within the bounds, randomize the trial order, and proceed with stimulation trials. The program waited at least 20 seconds between trials to allow the nerve to recover. In high amplitude or blocking techniques we increased the wait time up to 40 seconds. Recruitment curves consisted of 12-40 data points, with most trials using 20 points. We repeated these recruitment curves until all techniques were tested. Following all trials we euthanized the animal.

### Thresholds

For all animals and all experimental conditions, we determined the stimulation threshold to be the lowest current amplitude which produced motor activity over ± 25 µV. Our noise floor was typically less than ± 5 µV, and motor contractions were often just barely visible around ± 25 µV.

### Verification data

In some trials we want to compare recruitment data taken over the course of several hours, like the relationship between carrier frequency and threshold. In cases where we compared data that was not collected immediately sequentially, we also obtained a repetition of the first stimulation trial for verification to ensure the data was consistent. In order for a curve to be verified the threshold must not be more than 10% different than the initial value. Also, we wanted to ensure the relative recruitment of each muscle remained consistent. We calculated the mean squared error for each stimulation amplitude for each curve for both muscle groups. We normalized this value to the total area of the curve. We considered the data validated if this normalized error was less than 1. Examples of curves which pass and fail are shown in the supplement. All comparisons made in this paper are from data either taken immediately sequentially or with a passed verification trial.

### Data processing

Electromyogram (EMG) data was recorded using the hooked needles as electrodes. The signal was filtered using an AM Systems 1700 amplifier (second order Butterworth; passband = 100 – 500 Hz; gain = 100; 60 Hz notch) and digitized at a sample rate of 250,000 samples per second. Data was then digitally filtered with a 6th order Butterworth low pass filter at 650 Hz. We used MATLAB’s peak detect function to find all EMG peaks during the period of maximum stimulation amplitude (1 second for most trials). For recruitment curve data the EMG amplitude of the fifth largest peak is used, to accommodate outliers. If fewer than 5 peaks were identified, the recruitment value was set to zero.

### Mapping algorithm

To generate our estimated nerve map, we used the 72 trials from each mapping data set. For each given amplitude we used linear interpolation to estimate the EMG amplitude at each of the 6 stimulus locations for each muscle. These EMG amplitudes were used as the weights, and we estimated the location of the nerve branches based on a weighted average of the center of each stimulus region.

### Statistics and reproducibility

We perform no statistical tests; in most cases no test is appropriate. The data presented are gathered from 25 unique animals. We make a total of 45 claims (i.e., the sum of “n =“ statements) supported by these animals.

## Acknowledgements

This work was funded by NIH NS119390. Thanks to P. Iglesias and T. Meyer for manuscript feedback.

## Author Contributions

R.B.B and M.T.W conceived the study, designed and performed all experiments, conceived and designed all tools, methods, code, and techniques, analyzed data, generated figures, and wrote the manuscript. P.P.I. contributed to study conception, experimental design, and reviewed the manuscript.

