## Supplement for "Temporal interference current stimulation in peripheral nerves"

### Supplemental figures: Verification curves which pass and fail

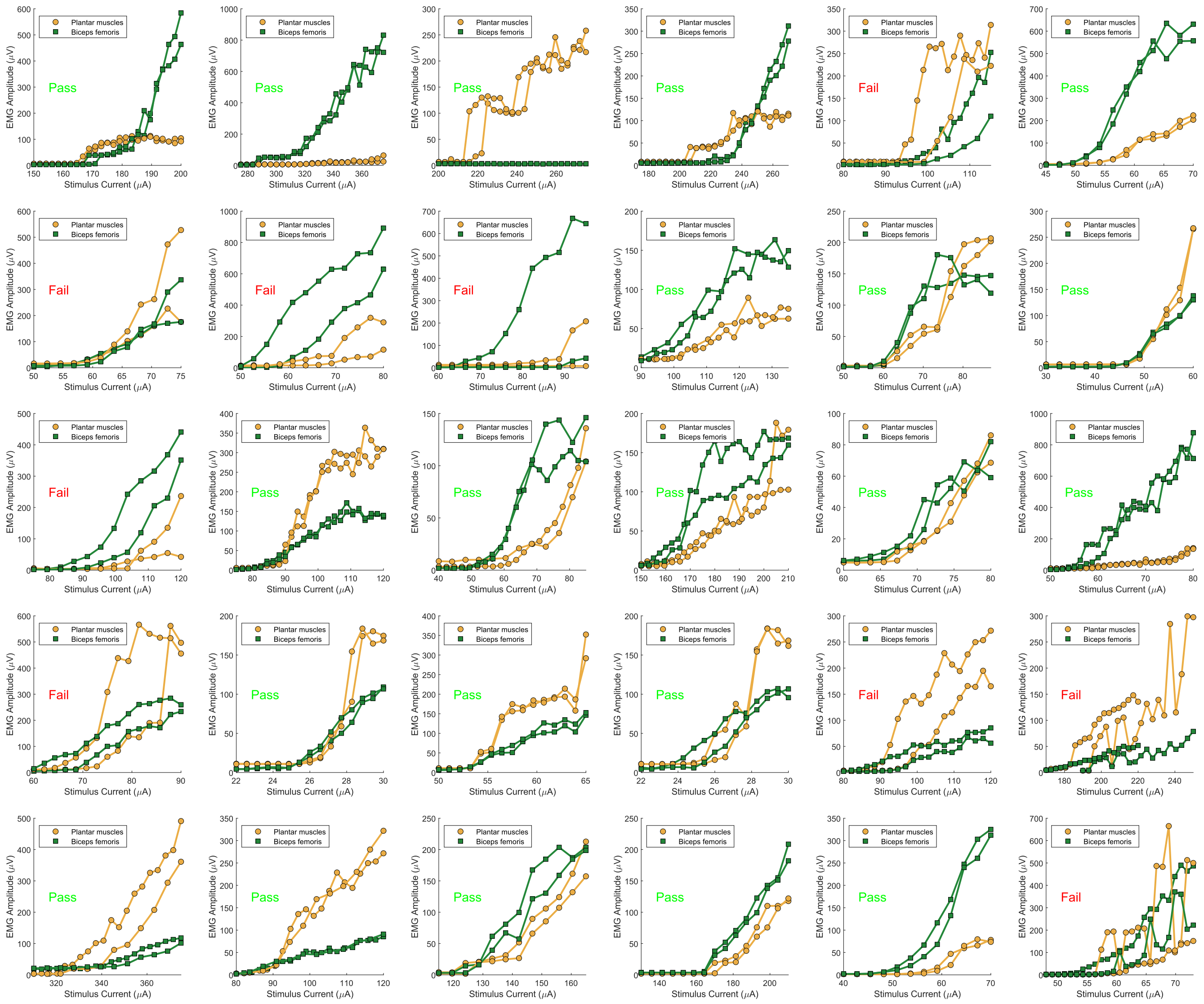


In each graph: A single strategy is tested. We then proceeded with an experiment and obtained data from many different strategies. To ensure comparisons between strategies are appropriate, we obtained a repetition of the first strategy tested. These trials are both plotted in each graph shown. The method for determination of verification is described in the Methods.

Circuit diagram:

We utilized a mirrored Howland current source as described in the Methods.

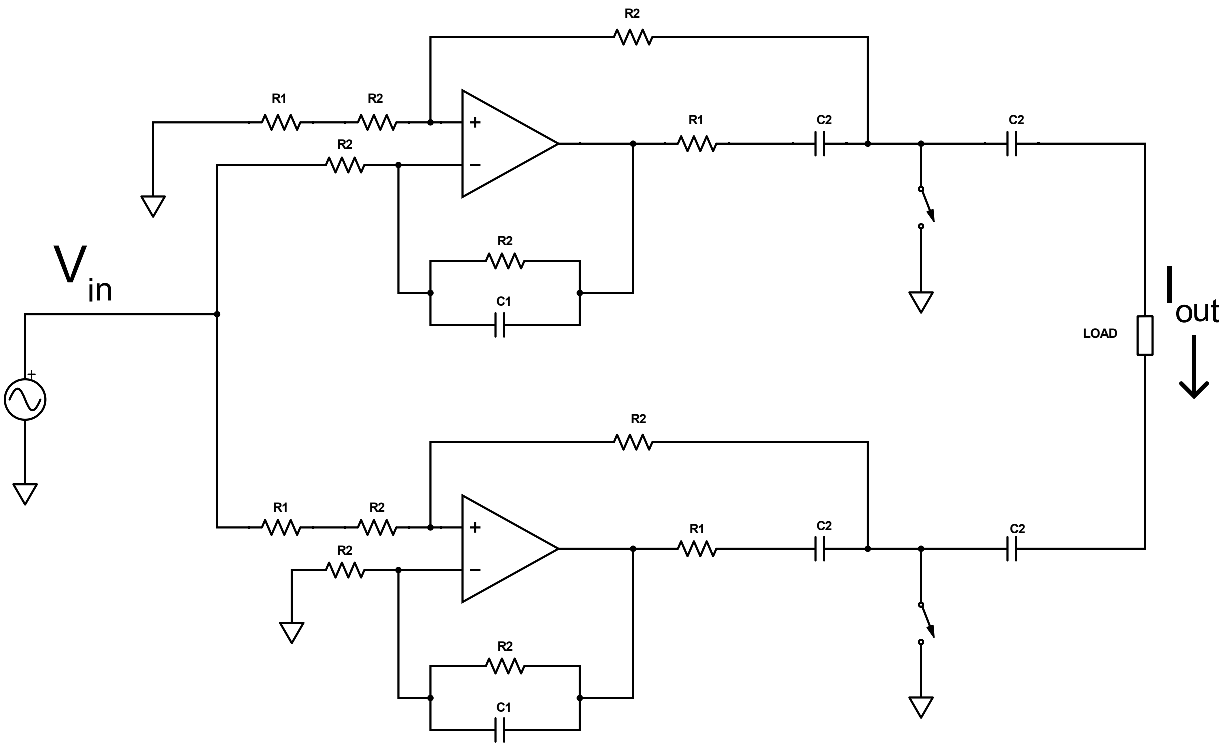


Values: $R_1$ = 660 Ω; $R_2$ = 100 kΩ; $C_1$ = 4.7 pF; $C_2$ = 2.2 µF
